# Mining all publicly available expression data to compute dynamic microbial transcriptional regulatory networks

**DOI:** 10.1101/2021.07.01.450581

**Authors:** Anand V. Sastry, Saugat Poudel, Kevin Rychel, Reo Yoo, Cameron R. Lamoureux, Siddharth Chauhan, Zachary B. Haiman, Tahani Al Bulushi, Yara Seif, Bernhard O. Palsson

**Affiliations:** Department of Bioengineering, University of California San Diego, La Jolla, CA, 92093, USA; Novo Nordisk Foundation Center for Biosustainability, Technical University of Denmark, Lyngby, 2800, Denmark

## Abstract

We are firmly in the era of biological big data. Millions of omics datasets are publicly accessible and can be employed to support scientific research or build a holistic view of an organism. Here, we introduce a workflow that converts all public gene expression data for a microbe into a dynamic representation of the organism’s transcriptional regulatory network. This five-step process walks researchers through the mining, processing, curation, analysis, and characterization of all available expression data, using *Bacillus subtilis* as an example. The resulting reconstruction of the *B. subtilis* regulatory network can be leveraged to predict new regulons and analyze datasets in the context of all published data. The results are hosted at https://imodulondb.org/, and additional analyses can be performed using the PyModulon Python package. As the number of publicly available datasets increases, this pipeline will be applicable to a wide range of microbial pathogens and cell factories.

## Introduction

Over the past few decades, advances in sequencing technologies have resulted in an exponential increase in the availability of public genomics datasets^1,2^. Public RNA sequencing datasets, in particular, have been integrated to provide a broad view of an organism’s transcriptomic state^3,4^, generate new biological hypotheses^5,6^, and infer co-expression networks and transcriptional regulation^7,8^.

Independent Component Analysis (ICA) has recently emerged as a promising method to extract knowledge from large transcriptomics compendia^9–16^. ICA is a machine learning algorithm designed to separate mixed signals into their original source components^17^. For example, given a set of microphones interspersed in a crowded room, ICA can be applied to the microphones’ recordings (with no additional input) to not only recreate the individual voices in the room, but also infer the relative distances between each microphone and each voice. Similarly, ICA can be applied to transcriptomics datasets to extract gene modules whose expression patterns are statistically independent to other genes. We call these independently modulated groups of genes *iModulons*. Most iModulons are nearly identical to regulons, or groups of genes regulated by the same transcriptional regulator, in model bacteria^9,10^, and can be used to discover new regulons or gene functions in less-characterized organisms^11,18^.

ICA simultaneously computes the activity of each iModulon under every experimental condition in the dataset, which represent the activity state of a transcriptional regulator. iModulon activities have intuitive interpretations, leading to a dynamic representation of an organism’s transcriptional regulatory network (TRN)^10,11,19–21^. Even for pairwise comparisons, iModulons can simplify the analysis from thousands of differentially expressed genes to merely dozens of differentially activated iModulons^22,23^.

iModulons have many properties that lend themselves to knowledge generation from large datasets. In terms of accuracy, ICA outcompeted 42 other regulatory module detection algorithms, including WGCNA and biclustering algorithms, in detecting known regulons across *E. coli*, yeast, and human transcriptomics data^24^. The FastICA algorithm^25^ is also relatively fast compared to deep learning approaches, computing components in a matter of minutes rather than hours^26^. Unlike many other decompositions, the independent components themselves are conserved across different datasets^27,28^, batches^29^ and dimensionalities within the same dataset^26,30^. This property enables us to use precomputed iModulon structures to interpret new transcriptomic datasets^9^. Altogether, these properties make ICA and iModulons a powerful tool to interpret the vast ocean of publicly available transcriptomic data to advance our understanding of transcriptome organization.

We have outlined a five-step workflow (**Figure 1a**) that enables researchers to build and characterize the iModulon structure for any microbe with sufficient public data. The first two steps are to download and process all publicly available RNA-seq data for a given organism. Third, the data must be inspected to ensure quality control, and curated to include all appropriate metadata. Next, ICA can be applied to the high-quality compendium to produce independent components. Finally, the independent components are processed into iModulons and can subsequently be characterized. To facilitate iModulon characterization, interpretation, and visualization, we present PyModulon, a Python library for iModulon analysis (https://pymodulon.readthedocs.io/en/latest/). We have made the entire workflow available on GitHub (https://github.com/avsastry/modulome-workflow), and will be disseminating iModulons through our interactive website (https://imodulondb.org/).

**Figure 1:**
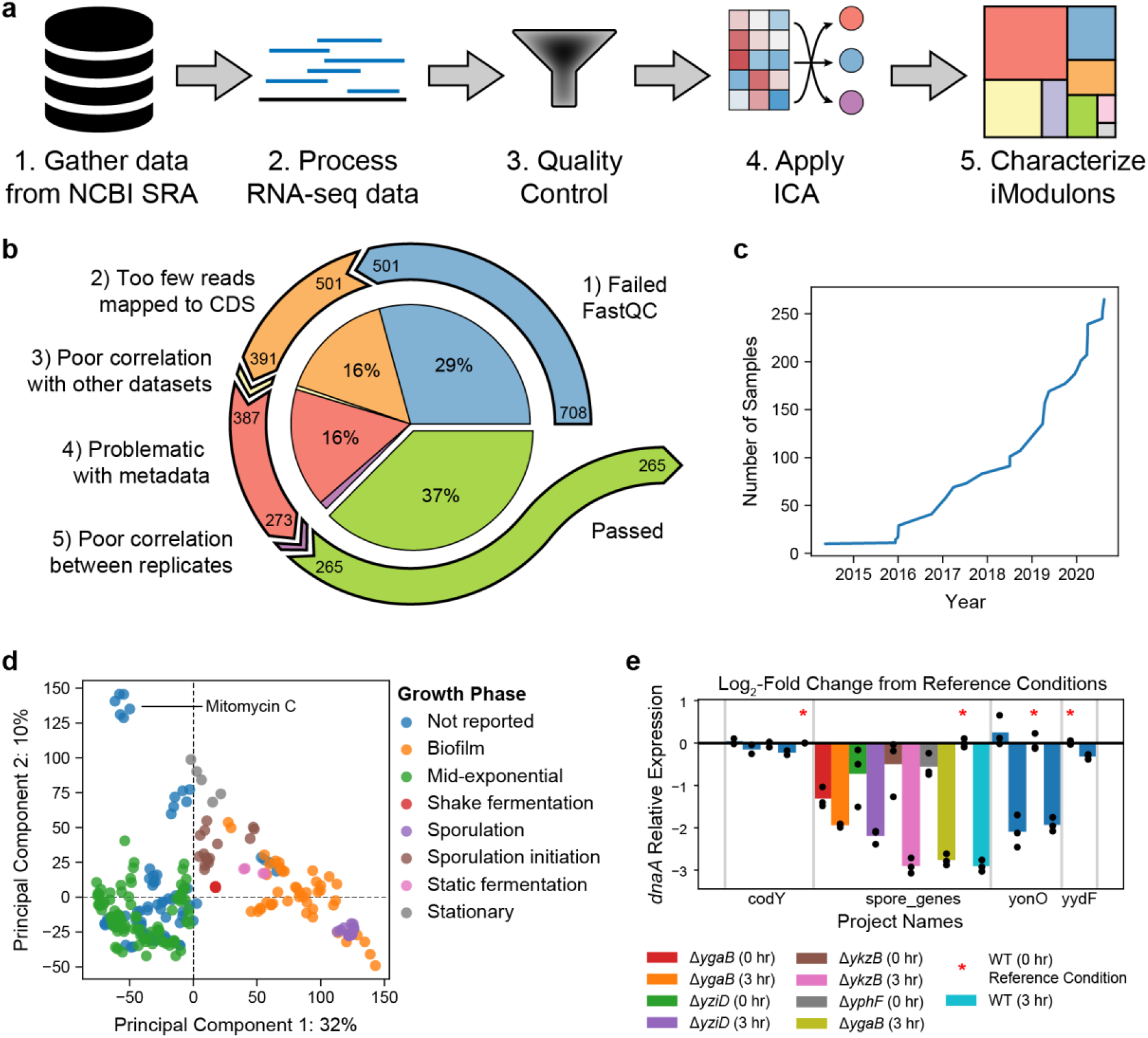
Overview of public *B. subtilis* RNA-seq data. **a)** Graphical representation of the five step workflow. **b)** Pie chart illustrating the quality control process. Numbers at the beginning of arrows represent the number of datasets before the quality control step, and numbers at the end represent the number of passed datasets after the step. **c)** Number of high-quality RNA-seq datasets for *B. subtilis* in NCBI SRA over time. **d)** Scatter plot of the top two principal components of the *B. subtilis* expression compendium. Points are colored based on the growth phase parsed from the literature. **e)** Bar chart showing the expression of *dnaA* across four projects. Points show individual replicates, while bars show the average expression for a given condition. Bars with a red star serve as the reference condition for the project. The legend describes the bars in the *spore_genes* project.

## Results

Here, we demonstrate how to build the iModulon structure of *Bacillus subtilis* from publicly available RNA-seq datasets using five steps (**Figure 1a**). All code to reproduce this pipeline is available at: https://github.com/avsastry/modulome-workflow/. Since this process results in the totality of iModulons that can currently be computed for an organism, we have named the resulting database as the “*B. subtilis* Modulome”.

### Steps 1 and 2: Compile and process all publicly available RNA-seq datasets for *B. subtilis*

Using Entrez Direct^31^, we have created a script that compiles the metadata for all publicly available RNA-seq data for a given organism in NCBI SRA (https://github.com/avsastry/modulome-workflow/tree/main/1_download_metadata). Since the iModulon structure improves with the number of datasets, we recommend that at least 50 unique conditions are available for an organism before proceeding with the remaining pipeline. As of August 2020, we identified 718 datasets labelled as *Bacillus subtilis* RNA-seq data.

Although iModulons can be computed from both microarray and RNA-seq datasets, microarray datasets tend to produce more uncharacterized iModulons, induce stronger batch effects through platform heterogeneity, and have been largely superseded by RNA-seq in current publications^27^. For these reasons, we have designed the first two steps specifically for compiling and processing RNA-seq data.

The *B. subtilis* datasets were subsequently processed using the RNA-seq pipeline available at https://github.com/avsastry/modulome-workflow/tree/main/2_process_data (**Figure S1**). Ten datasets failed to complete the processing pipeline, resulting in expression counts for 708 datasets.

### Step 3: Quality control, metadata curation, and normalization

The *B. subtilis* compendium was subjected to five quality control criteria (**Figure 1b**, https://github.com/avsastry/modulome-workflow/tree/main/3_quality_control). The final high-quality *B. subtilis* compendium contained 265 RNA-seq datasets (**Figure 1c**). As part of the quality control procedure, manual curation of experimental metadata was performed to identify which samples were replicates. Inaccurate or insufficient metadata reporting can hinder the widespread utilization of public data and potentially prevent subsequent interpretation^2,32^.

Therefore, we inspected the literature to identify the strain and media used in the study, any additional treatments or temperature changes, and the growth stage (e.g., mid-exponential, stationary, biofilm, etc.), if reported. During curation, we removed some non-traditional RNA-seq datasets, such as TermSeq or RiboSeq.

During this step, it is also recommended to compile a draft TRN, as it will be used to associate computed iModulons with known regulators. For model organisms, such as *B. subtilis* and *E. coli*, dedicated databases already exist with this information^33,34^. Although some resources are available for less-characterized organisms^35,36^, most information must be scraped from primary literature.

Although manual curation is the most time-consuming part of the workflow, it facilitates deep characterization of patterns in the gene expression compendium. For example, application of Principal Component Analysis (PCA) to the *B. subtilis* expression compendium revealed that a large portion of the expression variation could be explained by the growth stage (**Figure 1d**).

To obviate any batch effects resulting from combining different expression datasets, reference conditions are selected within each project to normalize each dataset. This ensures that nearly all independent components are due to biological variation, rather than technical variation. After normalization, gene expression and iModulon activities can only be compared within a project to a reference condition, rather than across projects (**Figure 1e**).

### Step 4: Run Independent Component Analysis

The *optICA* script (https://github.com/avsastry/modulome-workflow/tree/main/4_optICA) computes the optimal set of independent components and their activities (See **Supplementary Note 1**). To simplify analysis, we apply a threshold to each independent component (see Methods), resulting in gene sets called iModulons. This process resulted in 72 iModulons for the *B. subtilis* compendium that explained 67% of the expression variance in the compendium (**Figure S2**). Although the fraction of variance explained by the 72 iModulons is much lower than the fraction of variance explained by 72 principal components, explained variance has additional meaning in ICA compared to PCA. Principal components are a mathematical representation of the compendium and often lack biological interpretation. However, independent components and iModulons can be directly linked to transcriptional regulation (via the characterization in Step 5), revealing the true sources of expression variation in the compendium.

### Step 5: Characterize iModulons

To facilitate iModulon characterization, we developed the PyModulon Python package. Here, we will describe the contents of the package and how each module contributes to understanding information encoded in iModulons.

#### The IcaData Object

At the core of the PyModulon package is the *IcaData* object (**Figure 2a**). The object contains all data related to iModulons for a given dataset, including the **M** and **A** matrices (See **Supplementary Note 2**), the expression matrix, the thresholds used to define iModulons, and a scaffold TRN mined from literature. In addition, the IcaData object contains three annotation tables that store: (1) gene functions and annotations, (2) experimental parameters and metadata for samples, and (3) iModulon annotations and enrichments. The IcaData object is designed to facilitate gene set enrichment analysis, visualization of iModulon gene weights and activities, and comparisons of different iModulon structures. In addition, the object can be exported in JSON format for portability.

**Figure 2:**
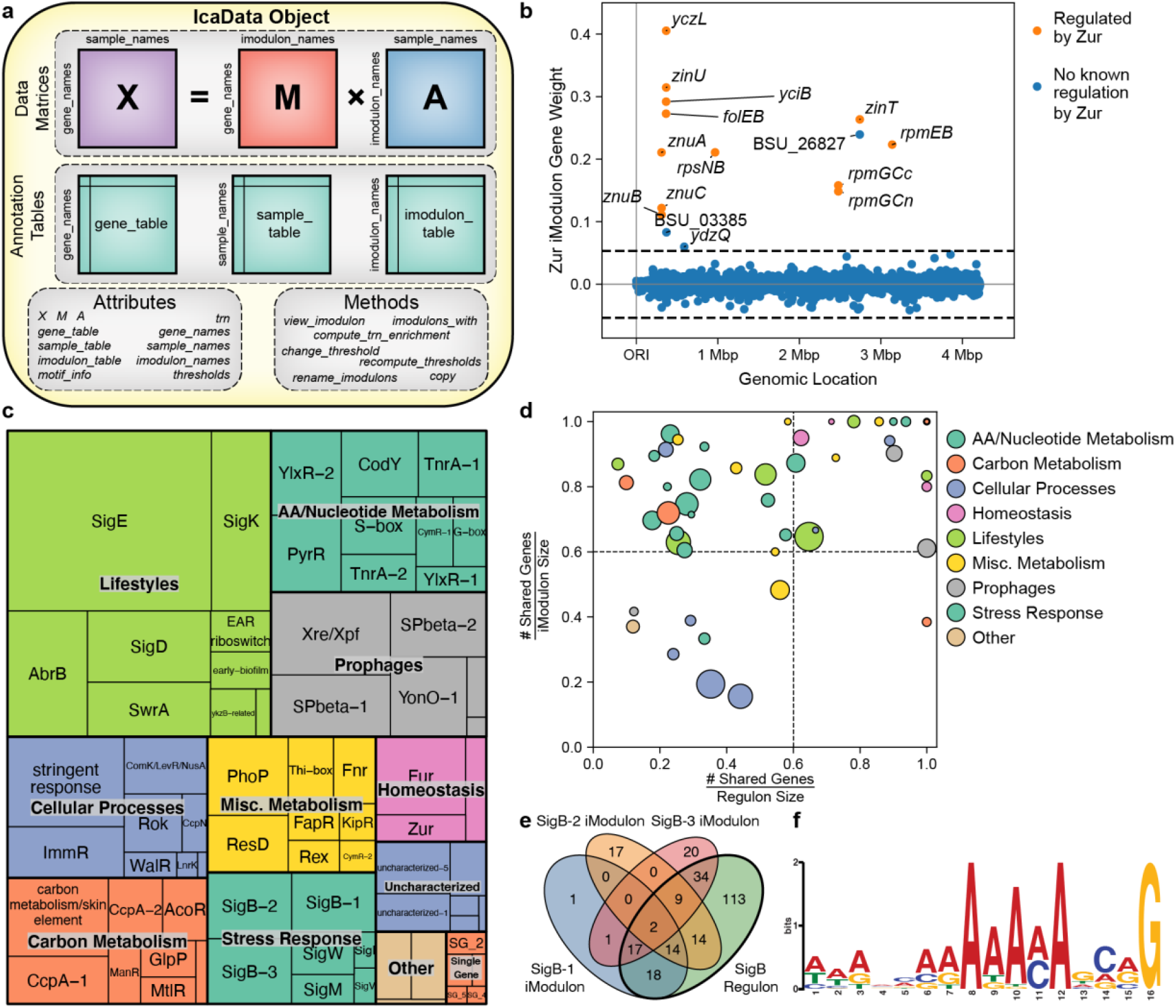
Overview of the *B. subtilis* iModulon structure. **(a)** Graphical representation of the *IcaData* object from PyModulon, illustrating the data, attributes, and methods stored in the object. **(b)** Example of an iModulon. Each point represents a gene. The x-axis shows the location of the gene in the genome, and the y-axis measures the weight of the gene in the Zur iModulon. Genes with prior evidence of Zur regulation are highlighted in orange. Genes outside the dashed black line are members of the Zur iModulon, whereas the genes inside the dashed black lines are not in the Zur iModulon. **(c)** Treemap of the 72 *B. subtilis* iModulons. The size of each box represents the fraction of expression variance that is explained by the iModulon. **(d)** Scatter plot comparing the overlap of each iModulon and its associated regulon(s). The circle size scales with the number of genes in the iModulon, and the color indicates the general category of the iModulon. **(e)** Venn diagram between the three SigB iModulons and the SigB regulon. **(f)** Motif identified upstream of all 58 genes in the ResD iModulon.

#### Determining iModulon Thresholds

Each independent component from ICA contains a gene weight for every gene in the genome. Only genes with weights above a specific threshold are considered to be in an iModulon. (**Figure 2b**). All thresholds are computed during initialization of the *IcaData* object using one of two algorithms: (1) the D’agostino method, which requires a moderately complete TRN, or (2) the K - means method, which can be used for less-characterized organisms (See Methods). Individual thresholds can be adjusted using the *change_threshold* method.

#### Characterizing iModulons

The *compute_trn_enrichments* method provides a user-friendly method to automatically identify iModulons that significantly overlap with regulons found in the literature. The method can be used to search for simple regulons (i.e., groups of genes regulated by a single regulator) or complex regulons (i.e., groups of genes regulated by a combination of regulators). This method is built on top of the *compute_annotation_enrichment* method, which can be used for gene set enrichment analysis against any gene set, such as gene ontology terms, KEGG pathways, plasmids, or phages.

Of the 72 *B. subtilis* iModulons, 52 iModulons represented the effects of known transcriptional regulators. Together, these *Regulatory* iModulons explain 57% of the variance in the dataset (**Figure 2c**). The iModulon recall and regulon recall can be used to assess the accuracy of regulator enrichments (**Figure 2d**). The iModulon recall is the fraction of the iModulon that is part of the pre-defined regulon from the literature, whereas the regulon recall is the fraction of the regulon that is captured by the iModulon. iModulons in the top left-quadrant often represent subsets of known regulons. For example, there are three iModulons that each capture different subsets of the SigB regulon (**Figure 2e**). Even though each iModulon contains a high fraction of genes regulated by SigB, none of the individual iModulons capture all of the genes thought to be regulated by SigB. Often, this separation represents independent regulatory modes for a single global regulator^22,23^. On the other hand, iModulons in the bottom right-quadrant provide evidence for new regulator binding sites. Even though only 28 of the 58 genes (48%) in the ResD iModulon have published ResD binding sites, we identified a conserved 16 base pair motif upstream of all 58 genes in the iModulon (**Figure 2f**).

Five additional iModulons were dominated by a single, high-coefficient gene, and are automatically identified by the method *find_single_gene_imodulons*. These *Single Gene (SG)* iModulons may arise from over-decomposition of the dataset^30,37^ or artificial knock-out or overexpression of single genes. Together, these iModulons contribute to 1% of the variance in the dataset.

The remaining 15 iModulons that could not be mapped to regulons derived from the literature present likely targets for the discovery of new regulons. Of those, the strongest candidates are the nine *Functional* iModulons, or iModulons that could be assigned a putative function. For example, one iModulon contains five genes in the same operon: *yvaC*, *yvaD*, *yvaE*, *yvaF*, and *azoRB*. Since YvaF is a putative transcription factor, we hypothesize that this iModulon is controlled by YvaF. Six *Uncharacterized* iModulons primarily contained either uncharacterized or unrelated genes, and contributed to 2% of the variance in the dataset.

Altogether, these 72 iModulons provide a quantitative framework for understanding the TRN of *B. subtilis*. This framework can be used to both re-interpret previously published studies in the context of the full compendium, and to rapidly analyze new data.

#### iModulon visualization

PyModulon contains a suite of functions to create publication-ready figures, as described here: https://pymodulon.readthedocs.io/en/latest/tutorials/plotting_functions.html. One such function computes a clustered heatmap of iModulon activities to identify correlated groups of iModulons (**Figure S3**). These iModulons often respond to a common stimulus, and represent a computational method to define stimulons^22,38^. Here, we guide readers through two stories that were discovered through iModulon visualizations.

First, we identified an uncharacterized iModulon that contains genes responsible for capsular polyglutamate synthesis, biofilm components, and synthesis of the peptide/polyketide antibiotic bacillaene^39^ (**Figure 3a**). This iModulon is activated in early biofilm production and stationary phase (**Figure 3b**), and the activities are correlated with the Since *B. subtilis* is a model organism for biofilm formation, and no single regulator is known to control all of these processes, this iModulon presents a compelling foundation for the potential discovery of a novel transcriptional regulator central to biofilm formation.

**Figure 3:**
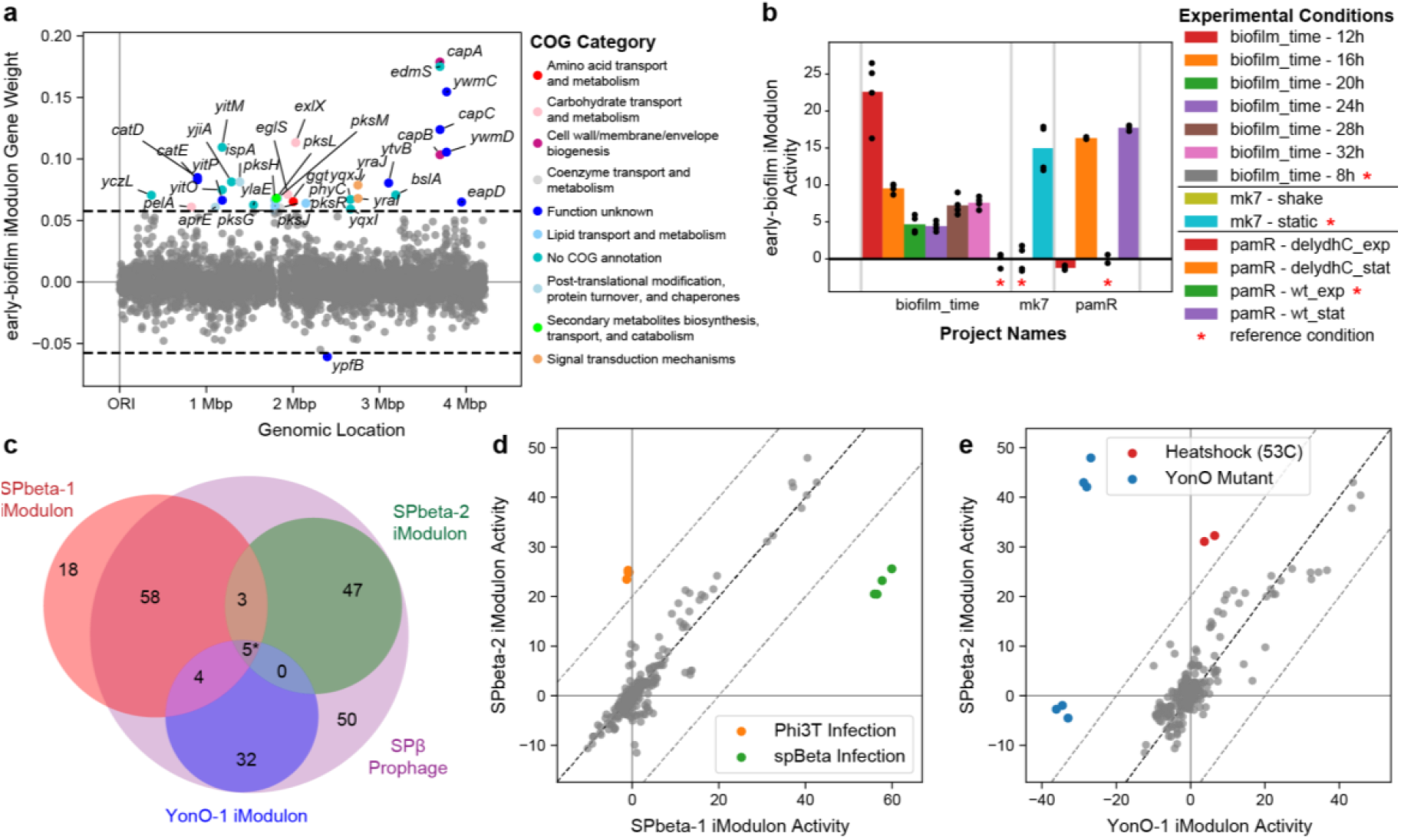
Examples of insights derived from iModulons. **(a)** Scatter plot of the gene weights in the newly discovered early-biofilm iModulon, created using the *plot_gene_weights* function. Genes outside the horizontal dashed black lines are in the iModulon, and genes are colored by their Cluster of Orthologous Gene (COG) category. **(b)** Bar plot of the iModulon activities for the early-biofilm iModulon, created from the *plot_activities* function. Individual points show iModulon activities for replicates, whereas the bars show the average activity for an experimental condition. Asterisks indicate the reference condition for each project. **(c)** Venn diagram comparing the SPbeta-1, SPbeta-2 and YonO-1 iModulons against the genes in the SPβ prophage. The asterisk indicates that one gene (*yozZ*) was in all three iModulons, but not in the prophage. **(d)** Scatter plot comparing the SPbeta-1 and SPbeta-2 iModulon activities, created from the *compare_activities* function. Each point represents a gene expression dataset under a specific condition. The center diagonal line is the 45-degree line of equal activities. **(e)** Scatter plot comparing the SPbeta-2 and YonO-1 iModulon activities. Each point represents a gene expression dataset under a specific condition.

In our second story, we gained a deeper understanding of the dynamics of a prophage’s expression profile by examining the interaction between iModulons. Three iModulons contain three distinct sections of the *B. subtilis* prophage SPβ (**Figure 3c**), one of which coincided with nearly all genes known to be transcribed by YonO, a recently discovered single subunit phage RNA polymerase^40^. All three iModulons shared 5 genes including the uncharacterized gene *yozZ*, which is not part of the SPβ prophage.

Although the activities of the three iModulons are nearly identical across most conditions, certain experimental conditions significantly deviate from the trend. The SPbeta-1 and SPbeta-2 iModulons diverge in a single experiment, where *B. subtilis* was infected with either the phage Phi3T or SPβ^41^ (**Figure 3d**). The SPbeta-1 iModulon has higher activity during infection with SPβ, whereas the SPbeta-2 iModulon has higher activity during infection with Phi3T.

The activities of the YonO-1 iModulon are nearly identical to the SPbeta-2 iModulon as well (**Figure 3e**). However, the two major differences include YonO mutant strains^40^ and a dataset where *B. subtilis* was exposed to heat shock at 53C^42^. The abnormally low YonO-1 iModulon activities are expected from the YonO mutant strain; however, the low activity during heat shock may indicate that the phage RNA polymerase YonO may be more sensitive to heat shock than the main *B. subtilis* RNA polymerase.

Every iModulons has a story similar to those described above. PyModulon visualization functions can help discover and illustrate these stories, accelerating generation of new hypotheses from re-analyzed data.

#### Estimating iModulon activities for external datasets

Once an iModulon structure has been defined for an organism, this structure can be inverted to infer iModulon activities for any new transcriptional dataset without re-running ICA (See Supplementary Text). This is implemented in the function *infer_activities*.

To demonstrate this functionality, we downloaded and processed an RNA-seq dataset tracking the development of *B. subtilis* biofilms^43^. This dataset was published after we compiled the transcriptomic compendium, so it was not used to compute the *B. subtilis* iModulon structure.

The resulting activity matrix (**Figure 4**) clearly shows that the sigma factors SigK and SigE are highly active during intermediate biofilm formation, between 2 days and 14 days, as compared to the reference condition of a fresh liquid culture. This likely explains the observation in the original study that the expression of transcription sigma factors increased during the intermediate biofilm stage^43^. The three iModulons related to the stress response sigma factor SigB exhibit an initial drop in activity but return to the reference levels after a day. iModulons related to carbon metabolism are generally down-regulated across the biofilm time-course, whereas iModulons encoding the PBSX phage (Xre/Xpf iModulon), capsular polysaccharide (EAR riboswitch), and competence (Rok) are up-regulated. Altogether, iModulons provide a fine-grained, while systems-level, view of the transcriptional changes occurring during biofilm formation.

**Figure 4:**
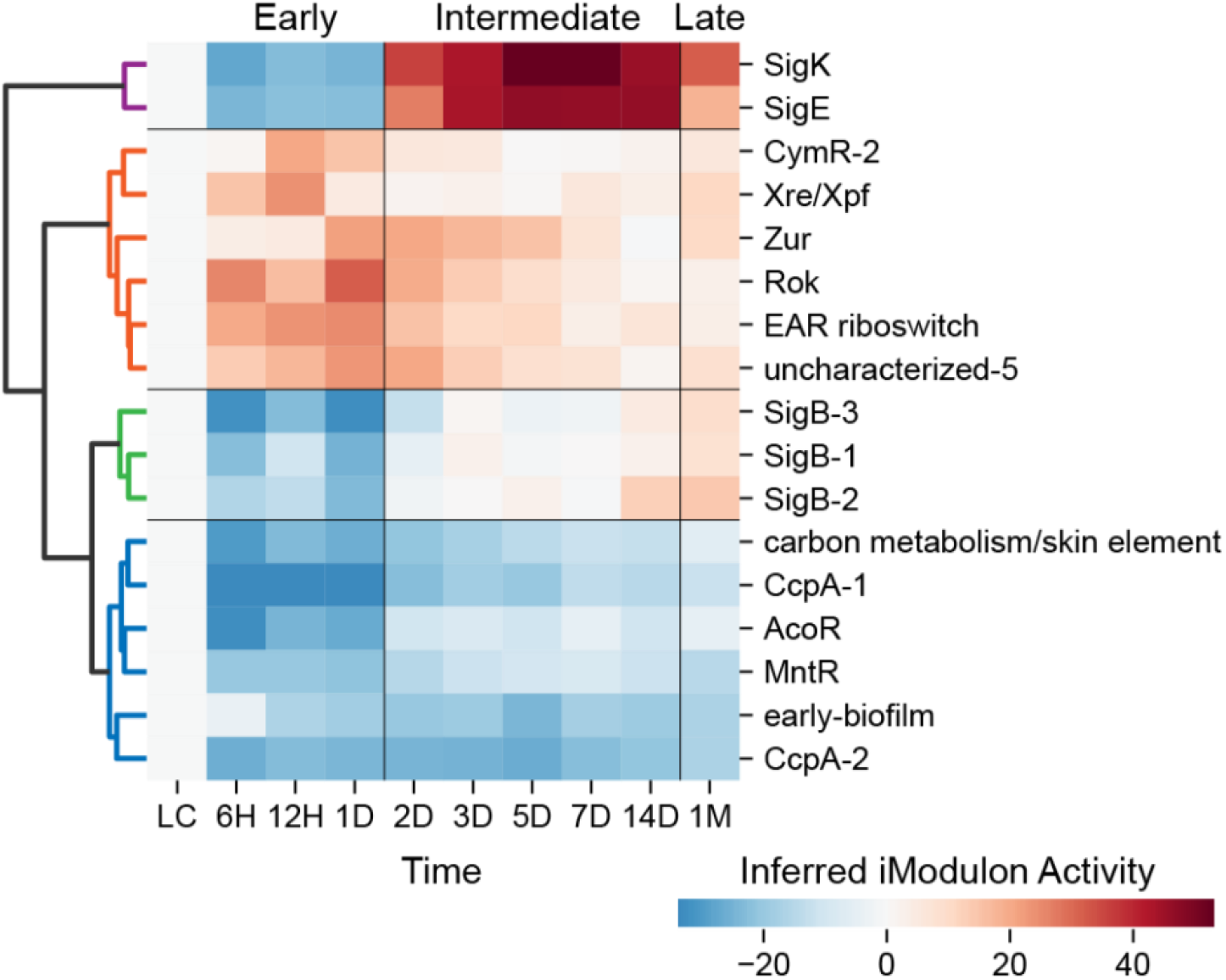
Clustermap of inferred iModulon activities across *B. subtilis* biofilm development. Abbreviations: LC - liquid culture, H - hours, D - days, M - month

#### Comparing iModulon structures

Previous studies have shown that similar iModulons can be found across disparate datasets^27,28^. To demonstrate this property, we use the *compare_ica* method to map the similarities between the iModulon structure presented here and an iModulon structure computed from a single microarray dataset^10,44^ (**Figure S4**). Of the 72 iModulons extracted from the RNA-seq compendium, 47 iModulons (65%) were highly similar to the microarray iModulons (**Figure 5a**). For example, nearly every gene in the Zur iModulon has nearly identical gene weights in both datasets (**Figure 5b**).

**Figure 5:**
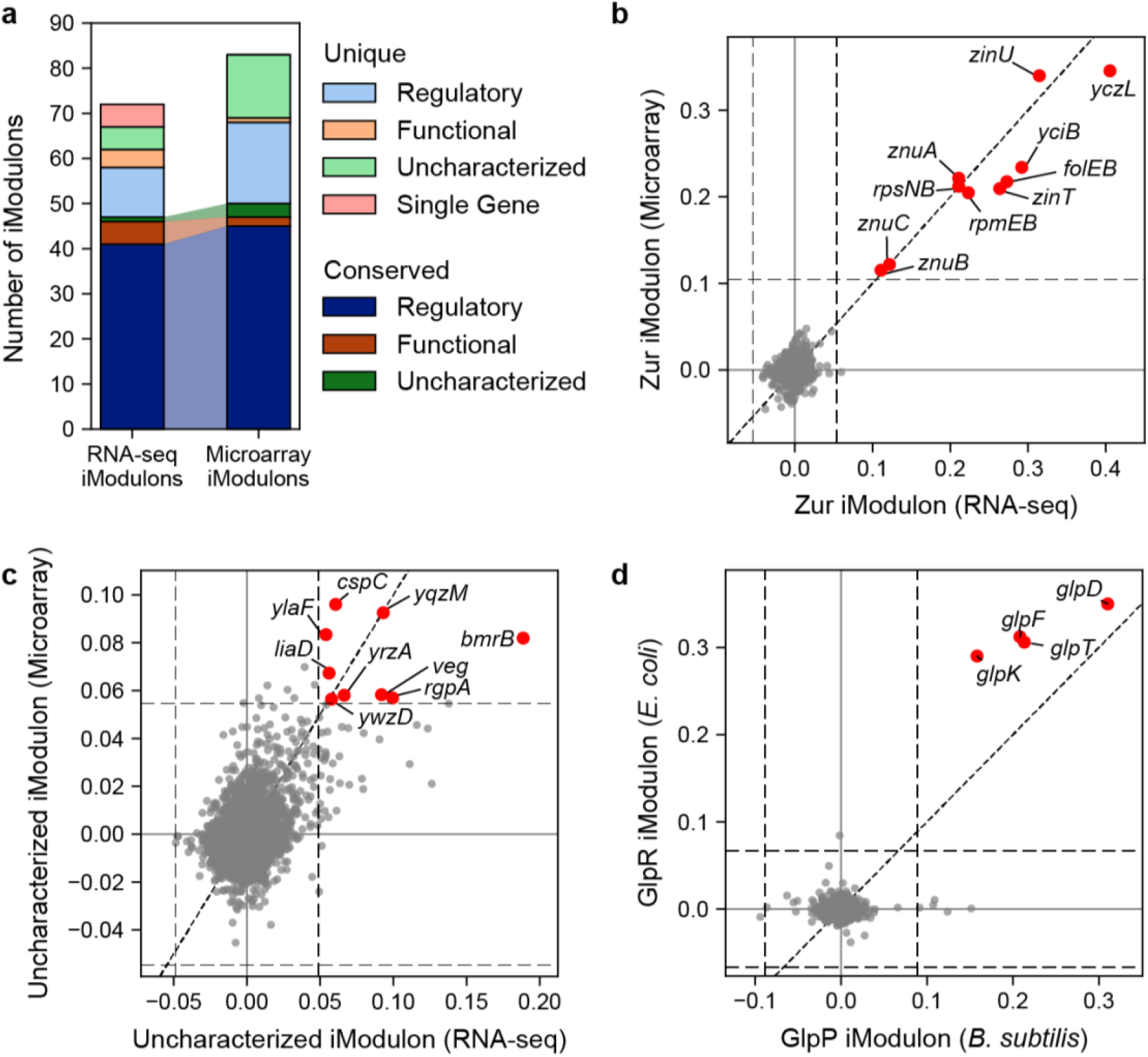
Comparison of iModulon structures. **(a)** Bar chart comparing the iModulons found in this RNA-seq dataset compared to a previous microarray dataset, colored by the type of iModulon. iModulons that are conserved between the two datasets are shown in a darker color. **(b-d)** Scatter plots comparing gene weights of iModulons found in different datasets, created using the *compare_gene_weights* function. Horizontal and vertical dashed lines indicate iModulon thresholds. Diagonal dashed line indicates the 45-degree line of equal gene weights. Genes in red are members of both iModulons. **(b)** Comparison of the Zur iModulon gene weights computed from the RNA-seq and microarray datasets. **(c)** Comparison of an uncharacterized iModulon found in both the RNA-seq and microarray datasets. **(d)** Comparison of the *B. subtilis* GlpP iModulon to the *E. coli* GlpR iModulon.

Presence of iModulons in two disparate datasets may indicate the existence of a previously unknown transcriptional regulator^27^. For example, an iModulon containing many uncharacterized genes was found in both datasets (**Figure 5c**). This is the same uncharacterized iModulon (uncharacterized-5) that was activated in the first few days of biofilm development, as shown in Figure 4. In the microarray dataset, this uncharacterized iModulon was downregulated in late sporulation. In total, this observation indicates that the genes in this iModulon are likely co-regulated by a transcriptional regulator that is related to biofilm development.

In addition, iModulons can be compared between organisms using gene orthology. We compared the *B. subtilis* iModulon structure to a previously published *E. coli* iModulon structure^9^, and found many orthologous iModulons (defined as iModulons containing orthologous genes with similar gene coefficients). We identified 22 iModulons in the *B. subtilis* dataset that were orthologous to *E. coli* iModulons (**Figure S5**). For example, the weights of the genes in the *B. subtilis* GlpP iModulon were nearly identical to their orthologs in the *E. coli* GlpR iModulon, indicating that these genes are modulated in similar ratios across the two organisms (**Figure 5d**).

#### Creating an iModulonDB web page

Since the iModulon decomposition for an organism is a knowledge-base that will be of interest to a broad audience of microbiologists, we created an interactive website (iModulonDB.org) for researchers to explore published iModulons^45^. iModulonDB lists all iModulons identified for a dataset, which link to interactive dashboards for each iModulon. Users can explore an iModulon’s member genes, activity levels across conditions, and comparisons with known regulons. Each gene also has its own page, with links to external databases, strengths of relationships to iModulons, and an expression profile. Within a dataset, users can search for genes or regulators of interest and browse all iModulons to obtain co-regulated gene sets or insights about condition-dependent activity. The results for the *Bacillus subtilis* dataset discussed here are available at https://iModulonDB.org/dataset.html?organism=b_subtilis&dataset=modulome.

The function *imodulondb_export* automatically generates all relevant files for an organism’s iModulonDB website from an *IcaData* object. Automated error checking and compatibility with the site javascript is ensured by the function *imodulondb_compatibility*. While these functions will work with a minimally curated *IcaData* object, the site experience is improved by the addition of hyperlinks for genes, transcription factors, metadata, and details about each sample. A Jupyter notebook^46^ guiding the addition of these annotations can be found at the following link (https://pymodulon.readthedocs.io/en/latest/tutorials/creating_an_imodulondb_dashboard.html).

Although the public iModulonDB site only contains our published decompositions, users can download its content from the github repository (https://github.com/SBRG/modulytics), add the files for their own decomposition, and then host a local version of the site with their data. Instructions for doing so are available in the iModulonDB github wiki (https://github.com/SBRG/modulytics/wiki/Adding-a-Private-Project). This method ensures privacy for new data while still enabling the search and dashboard utility of iModulonDB.

## Discussion

As of early 2021, the NCBI SRA database contains over 55,000 microbial RNA-seq datasets. We have provided a computational workflow to convert this mass of raw data into a transcriptomic knowledgebase for each organism that connects observed expression changes to their underlying transcriptional regulators. The resulting iModulon structures can accelerate the discovery of new regulons, aid in the characterization of genes with unknown function, and enhance the analysis of new datasets. In addition, the processed RNA-seq data is a resource for further studies of global expression patterns for the organism.

We demonstrated the potential of this workflow by analyzing all publicly available high-quality RNA-seq datasets for *B. subtilis*. Through iModulons, we identified potential new targets for known transcription factors, and new regulons that require further investigation to discover their transcriptional regulator. In addition, investigation into an iModulon induced by the deletion of YonO, a single subunit phage RNA polymerase, led to the observation that heat shock likely disrupts the protein’s function. The other iModulons capturing the SPβ prophage demonstrated that iModulons can provide understanding beyond just the TRN, and can be used to identify gene clusters such as prophages, pathogenicity islands, and biosynthetic gene clusters. Since the iModulons presented in this study represent the totality of expression signals that can be extracted from the currently available data, we have named this dataset the *B. subtilis* Modulome.

To assist further studies, we have developed three tools to compile, explore, and disseminate iModulons. First, we present a GitHub repository that walks through each analysis and figure discussed in this manuscript (https://github.com/avsastry/modulome-workflow). The pipeline is modular, as any step can be replaced with an alternative process, and the code in the repository can be modified for any new organism of interest. Second, we present PyModulon, a Python package for exploring iModulon properties, enrichments, and activities (https://pymodulon.readthedocs.io/en/latest/). Finally, all published iModulon structures can be uploaded to https://imodulondb.org, which contains interactive visualizations for any researcher to explore iModulons and global gene expression patterns.

There are currently many other microbes, including high-risk pathogens like *Mycobacterium tuberculosis*, with sufficient public expression data to compute iModulons. We foresee that this workflow will be broadly applied to all publicly available datasets, and any new large expression datasets, resulting in a Modulome for every available organism.

## Materials and Methods

### Compiling all public transcriptomics data for an organism

The NCBI Sequence Read Archive (SRA) is a public repository for sequencing data that is partnered with the EMBL European Nucleotide Archive (ENA), and the DNA Databank of Japan (DDBJ)^47^. We provide a script (https://github.com/avsastry/modulome-workflow/tree/main/download_metadata) that uses Entrez Direct^31^ to search for all public RNA-seq datasets and compile the metadata into a single tab-separated file. Each row in the file corresponds to a single experiment, and users may manually add private datasets.

### Processing prokaryotic RNA-seq data

The tab-separated metadata file can be directly piped into the prokaryotic RNA-seq processing pipeline (Figure 2a). This pipeline is implemented using Nextflow v20.01.0^48^ for reproducibility and scalability, and is available at https://github.com/avsastry/modulome-workflow. To process the complete *B. subtilis* RNA-seq dataset, we used Amazon Web Services (AWS) Batch to run the Nextflow pipeline. An alternative processing workflow for eukaryotic RNA-seq data is available in nf-core (https://nf-co.re/rnaseq)^49^.

The first step in the pipeline is to download the raw FASTQ files from NCBI using fasterq-dump (https://github.com/ncbi/sra-tools/wiki/HowTo:-fasterq-dump). Next, read trimming is performed using Trim Galore (https://www.bioinformatics.babraham.ac.uk/projects/trim_galore/) with the default options, followed by FastQC (http://www.bioinformatics.babraham.ac.uk/projects/fastqc/) on the trimmed reads. Next, reads are aligned to the genome using Bowtie^50^. The read direction is inferred using RSEQC^51^ before generating read counts using featureCounts^52^. Finally, all quality control metrics are compiled using MultiQC^53^ and the final expression dataset is reported in units of log-transformed Transcripts per Million (log-TPM).

### Quality Control and Data Normalization

To guarantee a high quality expression dataset for *B. subtilis*, data that failed any of the following four FASTQC metrics were discarded: per base sequence quality, per sequence quality scores, per base n content, and adapter content. Samples that contained under 500,000 reads mapped to coding sequences were also discarded. Hierarchical clustering was used to identify samples that did not conform to a typical expression profile, as these samples often use non-standard library preparation methods, such as ribosome sequencing and 3’ or 5’ end sequencing^3^.

Manual metadata curation was performed on the data that passed the first four quality control steps. Information including the strain description, base media, carbon source, treatments, and temperature were pulled from the literature. Each project was assigned a short unique name, and each condition within a project was also assigned a unique name to identify biological and technical replicates. After curation, samples were discarded if (a) metadata was not available, (b) samples did not have replicates, or (c) the Pearson R correlation between replicates was below 0.95. Finally, the log-TPM data within each project was centered to a project-specific reference condition.

### Computing the optimal number of robust Independent Components

To compute the optimal independent components, an extension of ICA was performed on the RNA-seq dataset as described in McConn et al.^37^.

Briefly, the scikit-learn (v0.23.2)^54^ implementation of FastICA^55^ was executed 100 times with random seeds and a convergence tolerance of 10^-7^. The resulting independent components (ICs) were clustered using DBSCAN^56^ to identify robust ICs, using an epsilon of 0.1 and minimum cluster seed size of 50. To account for identical with opposite signs, the following distance metric was used for computing the distance matrix:

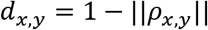

where *ρ_x,y_* is the Pearson correlation between components *x* and *y*. The final robust ICs were defined as the centroids of the cluster.

Since the number of dimensions selected in ICA can alter the results, we applied the above procedure to the *B. subtilis* dataset multiple times, ranging the number of dimensions from 10 to 260 (i.e., the approximate size of the dataset) with a step size of 10. To identify the optimal dimensionality, we compared the number of ICs with single genes to the number of ICs that were correlated (Pearson R > 0.7) with the ICs in the largest dimension (called “final components”). We selected the number of dimensions where the number of non-single gene ICs was equal to the number of final components in that dimension (**Figure S6**).

### Computing iModulon thresholds

Each independent component contains the contributions of each gene to the statistically independent source of variation. Most of these values are near zero for a given component. In order to identify the most significant genes in each component, we iteratively removed genes with the largest absolute value and computed the D’Agostino K2 test statistic^57^ for the resulting distribution. Once the test statistic dropped below a cutoff, we designated the removed genes as significant.

To identify this cutoff, we performed a sensitivity analysis on the concordance between significant genes in each component and known regulons. First, we isolated the 20 genes from each component with the highest absolute gene coefficients. We then compared each gene set against all known regulons using the two-sided Fisher’s exact test (FDR < 10^-5^). For each component with at least one significant enrichment, we selected the regulator with the lowest p-value.

Next, we varied the D’Agostino K2 test statistic from 50 through 2000 in increments of 50, and computed the F1-score (harmonic average between precision and recall) between each component and its linked regulator. The maximum value of the average F1-score across the components with linked regulators occurred at a test statistic of cutoff of 450 for the *B. subtilis* dataset.

For future datasets where a draft TRN is unavailable, an alternative method is proposed that is agnostic to regulator enrichments. The Sci-kit learn^54^ implementation of K-means clustering, using three clusters, can be applied to the absolute values of the gene weights in each independent component. All genes in the top two clusters are deemed significant genes in the iModulon.

### Compiling gene annotations

The gene annotation pipeline can be found at https://github.com/SBRG/pymodulon/blob/master/docs/tutorials/creating_the_gene_table.ipynb. Gene annotations were pulled from AL009126.3. Additionally, KEGG^58^ and Cluster of Orthologous Groups (COG) information were obtained using EggNOG mapper^59^. Uniprot IDs were obtained using the Uniprot ID mapper^60^, and operon information was obtained from Biocyc^61^. Gene ontology (GO) annotations were obtained from AmiGO2^62^. The known TRN was obtained from *Subti*Wiki^33^.

### Computing iModulon enrichments

iModulon enrichments against known regulons were computed using Fisher’s Exact Test, with the false discovery rate (FDR) controlled at 10^-5^ using the Benjamini-Hochberg correction. Fisher’s Exact Test was used to identify GO and KEGG annotations as well, with an FDR < 0.01.

Additional functions for gene set enrichment analysis are located in the *enrichment* module, including a generalized gene set enrichment function and an implementation of the Bonferroni-Hochberg false discovery rate (FDR).

### Motif search

Motif searches were performed using the *find_motif* function in PyModulon, which is a wrapper for MEME^63^. For each operon in an iModulon we searched for motifs in the sequence ranging from 500 base pairs upstream to 100 base pairs downstream of the gene start. For each iModulon, we searched for five motifs with zero or one occurrence per sequence, with an E-value threshold of 0.001. All motif widths were examined between 6 and 40 base pairs.

iModulon motifs were compared to known motifs using the *compare_motifs* function in PyModulon, which is a wrapper for TOMTOM^63^ using an E-value of 0.001. Motifs were compared against five prokaryotic motif databases^64–68^.

### Clustering iModulon activities

Global iModulon activity clustering was performed using the *clustermap* function in the Python Seaborn package^69^ using the following distance metric:

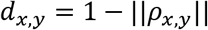

where ║*ρ_x,y_*║ is the absolute value of the Spearman R correlation between two iModulon activity profiles. The threshold for optimal clustering was determined by testing different distance thresholds to locate the maximum silhouette score (**Figure S3c**).

### Estimating iModulon activities for external datasets

The dataset for biofilm development was downloaded from NCBI GEO (GSE141305) and processed using the previously described pipeline. Data was centered on the reference condition (liquid culture). To infer the iModulon activities, the following equation was used:

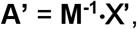

where **A’** is the inferred activity matrix, **M^-1^** is the pseudo-inverse of the precomputed independent components from the *B. subtilis* compendium, and **X’** is the expression matrix of the new data.

### Comparing iModulon structures

To compare iModulon structures between datasets from the same organism, each iModulon from one dataset was compared against every iModulon in the other dataset using the following distance metric:

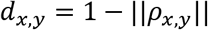

where ║*ρ_x,y_*║ is the absolute value of the Pearson R correlation between two independent components. iModulons with distances less than 0.25 were considered identical.

Before comparing iModulon structures across different organisms, gene orthology must be determined. As a simple metric, we used reciprocal BLAST hits to generate a one-to-one orthology between *E. coli* and *B. subtilis*. Once the orthologous pairs were determined, the iModulons were compared as described above.

## Acknowledgments

The authors would like to thank Dr. Amitesh Anand, Dr. Hyungyu Li, and Dr. Henrique Machado for informative discussions. This research used resources of the National Energy Research Scientific Computing Center, a DOE Office of Science User Facility supported by the Office of Science of the U.S. Department of Energy under Contract No. DE-AC02-05CH11231. This work was funded by the Novo Nordisk Foundation Center for Biosustainability and the Technical University of Denmark (grant number NNF10CC1016517).

