## Supplementary Information for "Mining all publicly available expression data to compute dynamic microbial transcriptional regulatory networks"

### Supporting information

#### Supplementary Notes

**Supplementary Note 1:**
FastICA is one of the most popular algorithms for ICA [1]. However, there are two considerations when running the FastICA algorithm. First, FastICA is a stochastic algorithm that results in different sets of independent components during different runs. Several solutions exist to identify components that are robust to random initialization [2,3]. In addition, one must set the number of components for ICA *a priori*. To address these issues, we previously published the *optICA* extension of the FastICA algorithm to identify robust components at the optimal dimensionality [4].

Briefly, *optICA* performs FastICA 100 times at a range of different dimensions. To identify the robust components at each dimension, the components are clustered to identify which components exist in at least 50% of the runs. This process is repeated across a range of specified dimensions to track the number of robust components present at each dimension (**Figure S6**). To detect under-decomposition, we also track the number of components at each dimension that are similar to the components found in the largest possible dimension (i.e. “final” components). To detect over-decomposition, we track the number of components that are dominated by a single gene (i.e. “single-gene” components). The optimal dimensionality exists where the number of “final” components overtakes the number of non-single-gene components.

**Supplementary Note 2:**
ICA is a matrix decomposition algorithm that takes a data matrix (**X**) and decomposes it into two smaller matrices (**M** and **A**), where **X** = **MA**. For this study, the **X** matrix contains a collection of gene expression profiles, where each column is a gene expression profile under a specific condition, and each row represents the expression of a single gene. Each column of the **M** matrix contains an independent component. Each independent component contains a weighting for each gene, and most gene weights in a component are near zero (**Figure 2b**). To identify genes with significant weightings, we apply a unique threshold to each component (See Methods), and consider genes with weightings outside this threshold as part of the iModulon.

Each column of the **A** matrix contains the iModulon activities for a specific expression profile. Since we centered each expression profile to a project-specific reference condition, we can only compare activities across a project, rather than between projects. On the other hand, the gene weights represent the strength of regulation for a transcription factor on each gene. A gene with a high weight for a specific iModulon would be more sensitive to changes in iModulon activity than a gene with a low weight. Overall, the relative expression of a single gene under a specific condition is the sum of gene weights across all iModulons, each weighted by the conditions-specific iModulon activity (i.e. the product of the **M** and **A** matrices results in the expression matrix (**X**)).

### Supporting Figures

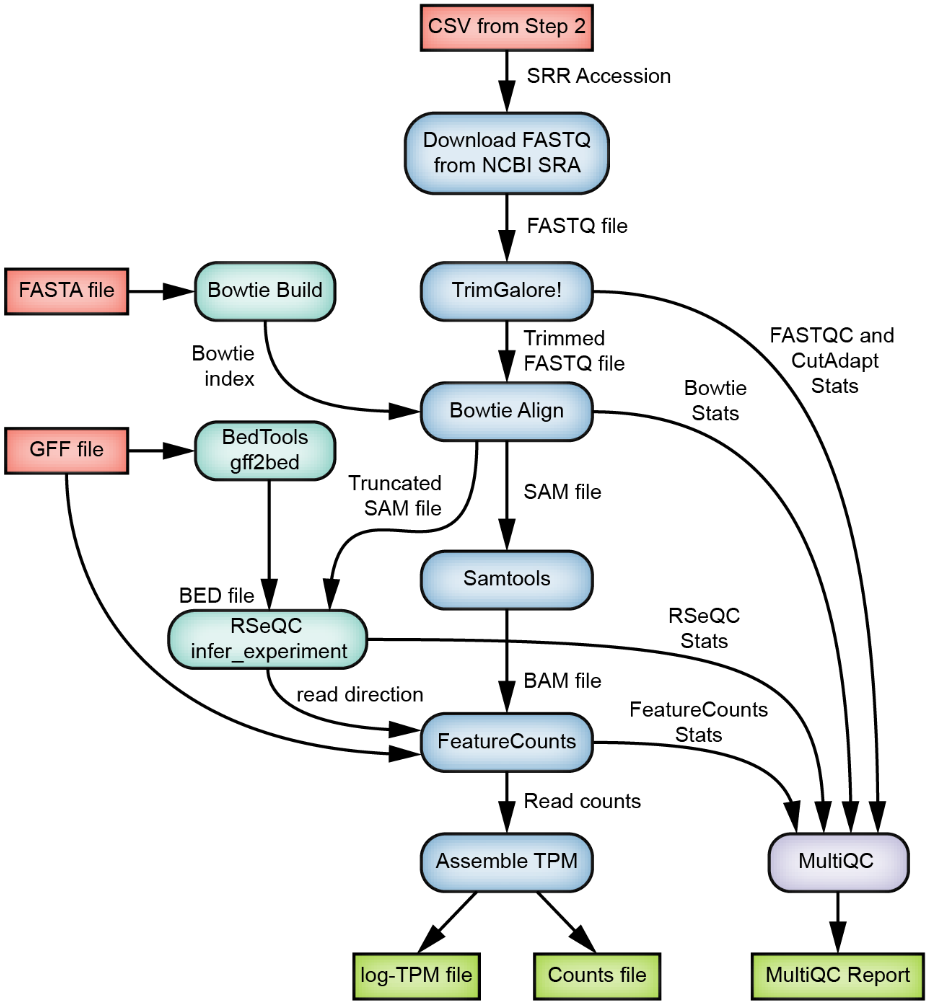


**Figure S1:** Processing pipeline for RNA-seq datasets. Input files are boxed in red, and output files are boxed in green.

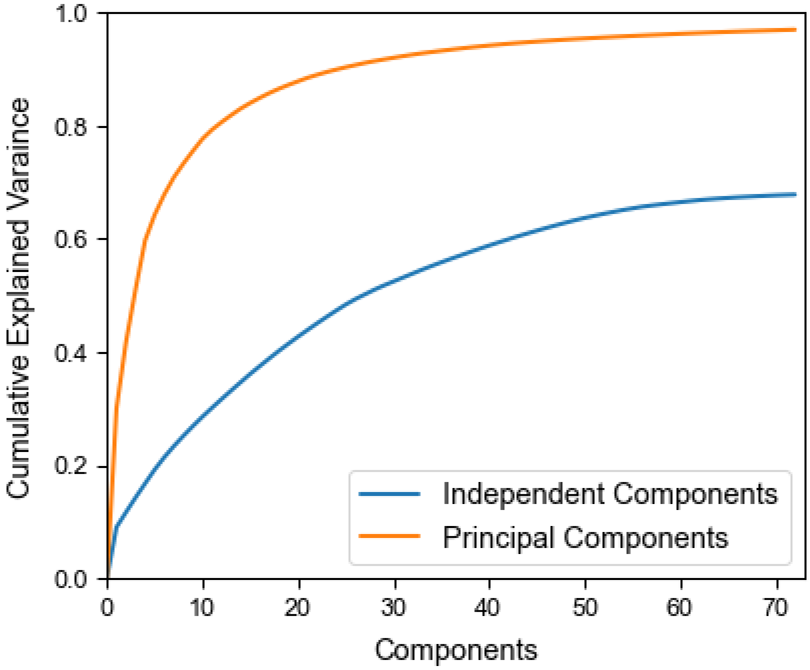


**Figure S2:** Cumulative fraction of expression variance explained by each component for Principal Component Analysis and Independent Component Analysis.

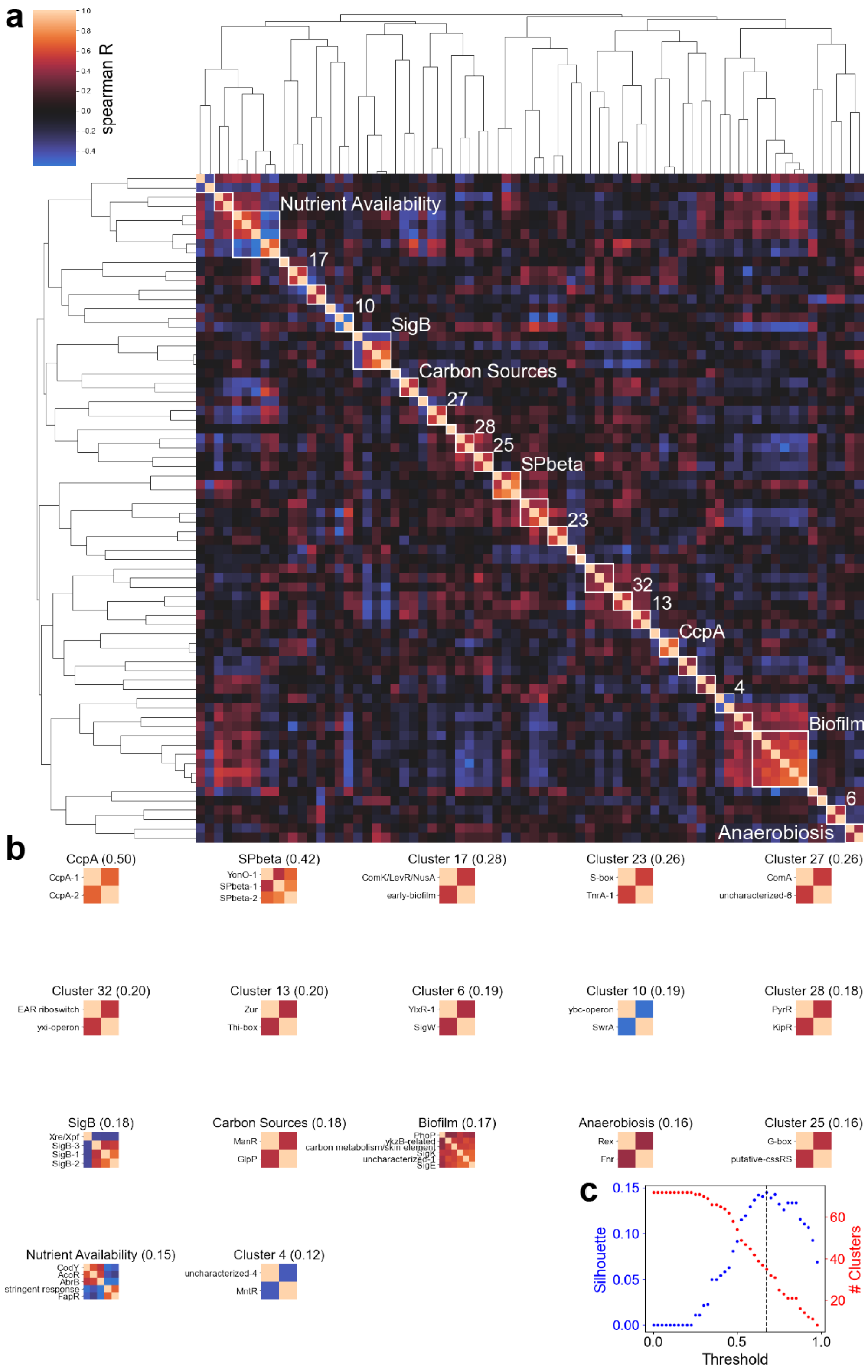


**Figure S3:** (a) Clustered heatmap of Spearman correlations between iModulon activities. (b) Names of iModulons in the best clusters. (c) Sensitivity analysis to identify the clustering threshold.

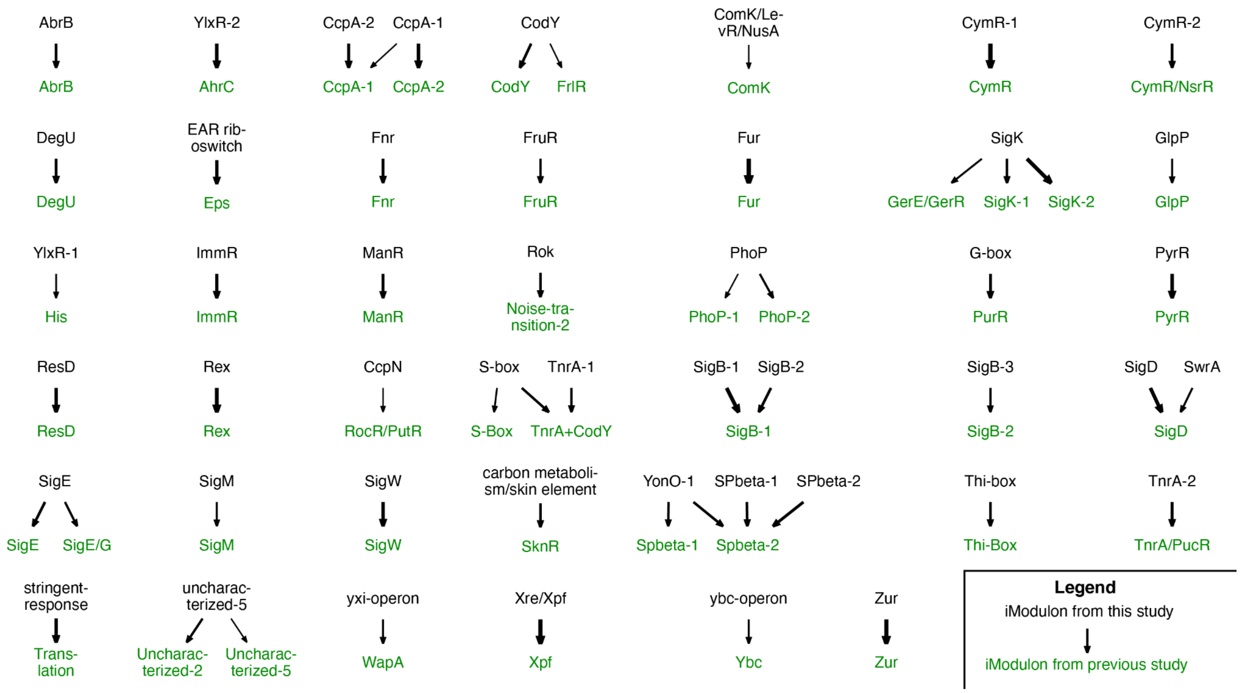


**Figure S4:** Comparison of iModulons computed from the compendium presented in this study (black) against iModulons computed from a microarray dataset (green). Arrow width indicates the Pearson R correlation between the components.

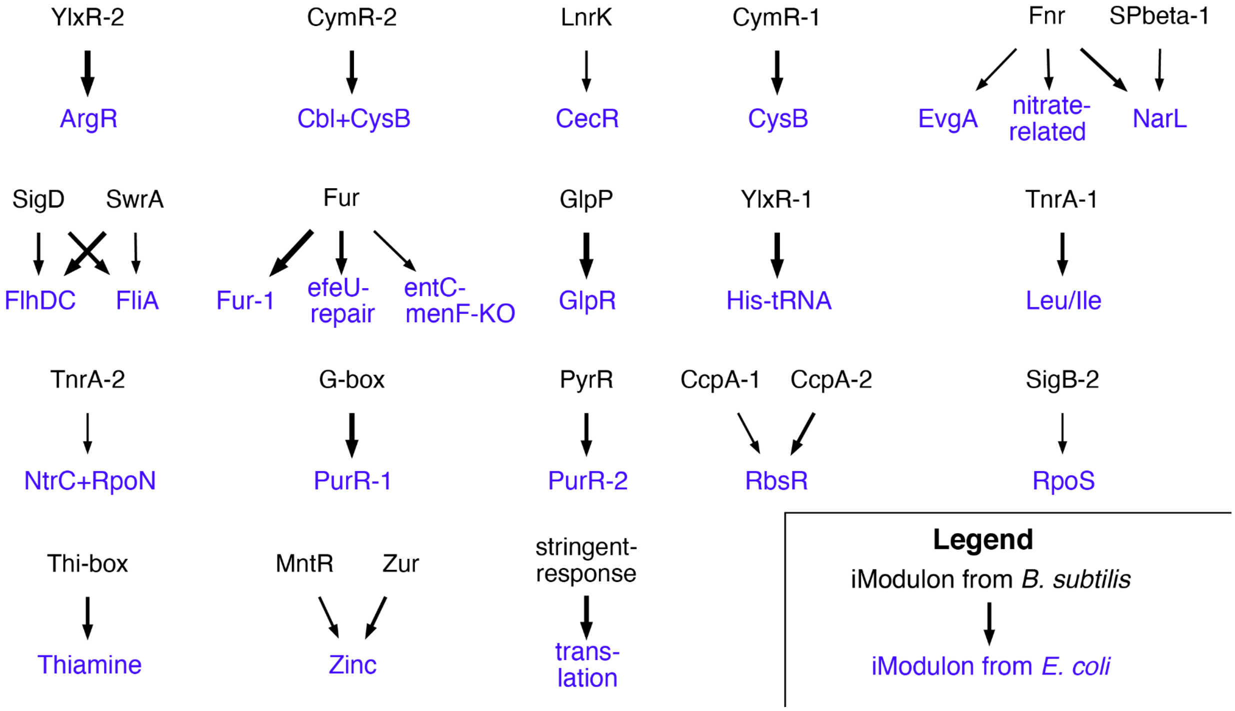


**Figure S5:** Comparison of iModulons computed from the *B. subtilis* RNA-seq compendium presented in this study (black) against iModulons computed from an *E. coli* RNA-seq compendium (blue). Arrow width indicates the Pearson R correlation between the components.

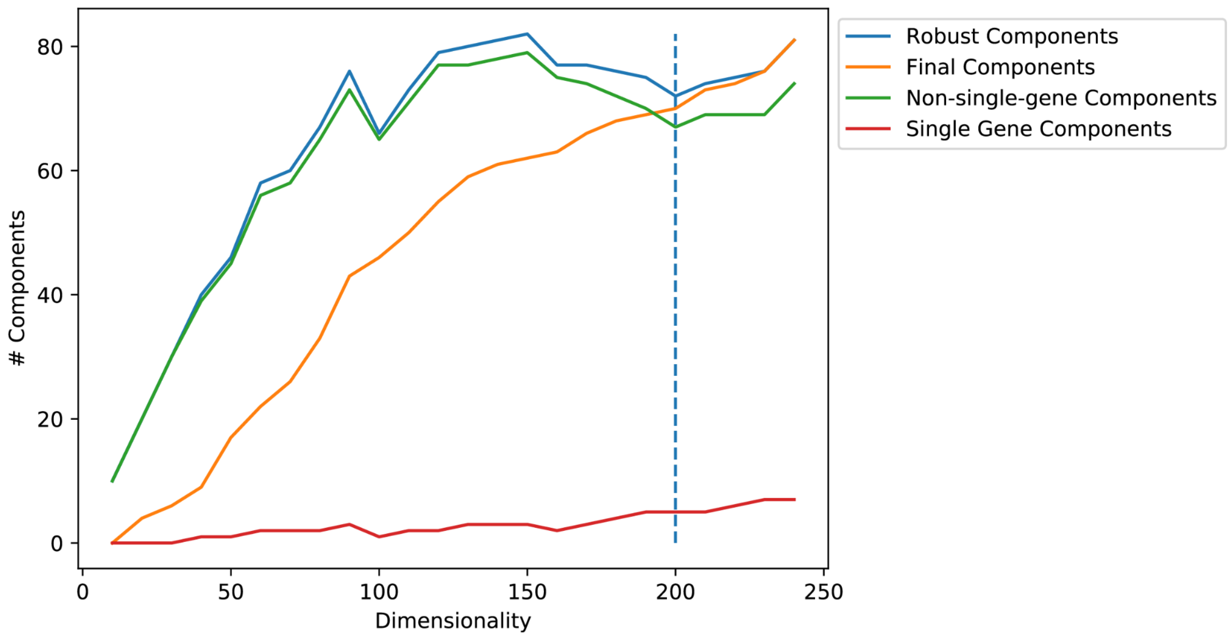


**Figure S6:** Number of components computed at each dimensionality. Robust components are components that are found in over 50% of randomly initialized runs of ICA. Final components are components that have an absolute Pearson correlation coefficient greater than 0.7 with the set of components found at the highest dimension. Single-gene components are components where a single gene dominates the gene coefficients, and non-single-gene components are all components that are not single gene components. The optimal dimensionality is found when the number of non-single-gene components is surpassed by the final components. See **Supplementary Note 1** and McConn et al. [4] for more information.
